# Gfral-expressing Neurons Suppress Food Intake via Aversive Pathways

**DOI:** 10.1101/2020.05.11.088773

**Authors:** Paul V. Sabatini, Henriette Frikke-Schmidt, Joe Arthurs, Desiree Gordian, Anita Patel, Jessica M. Adams, Jine Wang, Sebastien Beck Jørgensen, David P. Olson, Richard D. Palmiter, Martin G Myers, Randy J. Seeley

## Abstract

To determine the function and mechanisms of action for hindbrain neurons that express GFRAL, the receptor for the anorexigenic peptide, GDF-15, we generated *Gfral^cre^* and conditional *Gfral^CreERT^* mice. While signals of infection or pathophysiologic states (rather than meal ingestion) stimulate GFRAL neurons, the artificial activation of *Gfral^Cre^*- expressing neurons inhibited feeding, decreased gastric emptying, and promoted a conditioned taste aversion (CTA). Additionally, activation of the smaller population of GFRAL neurons captured by the *Gfral^CreERT^* allele decreased gastric emptying and produced a CTA without suppressing food intake, suggesting that GFRAL neurons primarily modulate gastric physiology and stimulate aversive responses. GFRAL neurons most strongly innervated the parabrachial nucleus (PBN), where they targeted CGRP-expressing (CGRP^PBN^) neurons. Silencing CGRP^PBN^ neurons abrogated the aversive and anorexic effects of GDF-15. These findings suggest that GFRAL neurons link non-meal-associated, pathophysiologic signals to the aversive suppression of nutrient uptake and absorption.

## Introduction

Several lines of evidence link high circulating levels of the cytokine, growth and development factor 15 (GDF-15; also called MIC-1), to the reduced hunger, decreased food intake, and weight loss that occur in a variety of (Johnen et al., 2007). Consequently, many have suggested that GDF-15 and its analogues could be used to decrease feeding and promote weight loss in individuals with obesity. Indeed, exogenous GDF-15 treatment leads to substantial weight loss in mice, rats and non-human primates (Emmerson et al., 2017; Hsu et al., 2017; Mullican et al., 2017; Yang et al., 2017).

The identification of the receptor that mediates the anorexigenic effects of GDF-15, GDNF family receptor α–like (GFRAL), has provided an opportunity to understand novel mechanisms of food-intake suppression. While a wide range of tissues synthesize and secrete GDF-15 in response to cellular stress (Coll et al., 2019; Montero et al., 2016; Nakayasu et al., 2020; Patel et al., 2019), GFRAL has a uniquely narrow expression pattern restricted to a small population of CNS neurons. Importantly, the hypothalamus and other forebrain regions contain no GFRAL, but rather GFRAL-expressing cells reside in the hindbrain, primarily in the area postrema (AP), with a smaller number in the adjacent *nucleus tractus solitarius* (NTS)(Emmerson et al., 2017; Hsu et al., 2017; Mullican et al., 2017; Yang et al., 2017).

Because much of the machinery that controls long-term feeding and body weight depends on hypothalamic circuits (and many current pharmacological strategies to produce safe and efficacious weight loss engage these hypothalamic circuits), it is somewhat surprising that a very small population of neurons in the AP might mediate the effects of GDF-15 on food intake and body weight. Indeed, the AP has received considerably less attention as a target for producing weight loss. The neurons targeted by GDF-15 can modulate body weight, however, and understanding their function and the downstream circuits by which they act will be crucial for the advancement of GDF-15-based therapeutics. To that end, we developed mice that express constitutive or inducible Cre recombinase in GFRAL-expressing neurons. This allowed us to use Cre-dependent systems to study and manipulate GFRAL neurons, revealing their regulation, function, and downstream mediators.

## Star Methods

### Lead Contact and Materials and availability

Further information and requests for resources and reagents should be directed to and will be fulfilled by the Lead Contact, Randy Seeley

### Experimental model and subject details

Mice were bred in the Unit for Laboratory Animal Medicine at the University of Michigan. These mice and the procedures performed were approved by the University of Michigan Committee on the Use and Care of Animals and in accordance with Association for the Assessment and Approval of Laboratory Animal Care and National Institutes of Health guidelines. Mice were provided with *ad libitum* access to food (Purina Lab Diet 5001, unless otherwise specified) and water (except as noted below) in temperature-controlled (25°C) rooms on a 12 h light-dark cycle with daily health status checks.

*Gfral^Cre^* and *Gfral^CreERT^* animals were generated with dual gRNA (aUgUcUgUgUgacUaaccaa & GUaaaaUgUgacaaaUUUgc) and plasmid donors through direct blastocyst nuclear injection performed by the University of Michigan Transgenic core. Potential founders were screened with generic Cre primers (TGAGGTTCGCAAGAACCTGATGGA Cre F; AGGGCGCGAGTTGATAGCT Cre R); mice that were genotyped as Cre+ were subsequently screened for targeted incorporation (5’: ATA TTG CTG CCC CCA GAA GC *Gfral* WT F; CAC ATT GCC AAA AGA CGG CA *Gfral-IRES-Cre*(*ERT*) R; 3’ Gca tga tga atc tgc agg gag ag *Gfral-IRES-CreERT* F; ACC AGC CAG CTA TCA ACT CG *Gfral-IRES-Cre* F; TTG GAG GGA ACA TTG CGA CA *Gfral* WT R). For genotyping *Gfral^Cre^* and *Gfral^CreERT^* mice, standard PCR was performed on DNA isolated from tail clips (Genotyping primers: GAG ATC CCT CAT CCA TCG AAA TAG Gfral WT F; CAC ATT GCC AAA AGA CGG CA Gfral Cre/CreERT R; CTG CAC ACA CTA ATT ATG ATT AGT TCC Gfral WT R). These lines were crossed to *Rosa26*^eGFP-L10a^ (a generous gift from D.Olson), hM3Dq^Tg^ (JAX strain 026220), and *Rosa26^S^y^nTdT^* (JAX strain 012570).

For studies performed at the University of Washington, all experiments were approved by the University of Washington Institutional Animal Care and Use Committee and were performed in accordance with the guidelines described in the U.S. National Institutes of Health Guide for the Care and Use of Laboratory Animals. *Calca^Cre:GFP^* mice were described (Carter et al., 2013).

## Method Details

### Stereotaxic surgeries and infusion site verification in rats

Male Long Evans rats were fed Tso’s 40% butter-fat diet for up to 8 weeks. Two weeks prior to surgery they were singly housed and handled daily. On the day of surgery, each rat was anesthetized using 5% isoflurane and maintained on 2% isoflurane. 0.03 mg/kg buprenex and 0.5 mg/kg meloxicam were provided as analgesia. Each rat was placed in the stereotaxic frame and an incision was made through the skin across the skull. The bregma and the lambda were both identified and the skull was adjusted to be horizontal between these. For AP cannulation a hole was drilled on the ridge at the caudal end of the skull (approximately 15 mm caudal to bregma and 0 mm lateral to the midline. The bregma distance varied between rats depending of the location of the skull ridge). For

NTS cannulation a hole was drilled 0.5 mm rostral to the skull ridge and 0.9 mm lateral to the midline. Cannulation depth for both AP and NTS, respectively, varied between 9.4 and 10.2 mm ventral to bregma. The exact depth was tested in 3 to 5 rats in each cohort prior to the actual surgery as skull thickness and size varied between high-fat fed cohorts. For all cannulation surgeries, the cannula was held in place by dental acrylic (fujicem) and the rostral part of the incision was closed with wound clips. Animals received analgesia for 2 days after surgery: 0.03 mg/kg buprenex twice a day and 0.5 mg/kg ostilox once daily. Wound clips were removed 7 days post-surgery.

At the end of each experiment the rats were euthanized using CO_2_ and decapitation. 0.5 μl blue dye was then slowly infused over 2 minute and the brain was extracted and frozen in tissue tech on dry ice. Each brain was sectioned on a cryostat and injection site was verified by the presence of the blue dye using a rat brain atlas (Paxinos and Watson, 2013) for reference.

### Neuropile infusions and food intake in rats

After surgery, the rats were handled daily with the dummy cannula being pulled out and inserted to habituate rats to cannula manipulations. Infusions started 2 weeks after surgery, when all rats were fully healed. On each infusion day, food was removed 2 hours prior to lights out. One hour later infusions were initiated with 1 μl of vehicle, GDF-15 (doses of 0.33, 1, or 2 μg), or salmon calcitonin (0.4 μg) being slowly infused over 1 minute with an additional one-minute wait before the infusion needle was removed.

Food was then given back exactly 1 hour after the infusion. Food intake was measured 1,2, and 23 hours after infusion. Each rat had 3 infusions with doses randomized for infusion day and were allowed a minimum of 1 week of recovery between each infusion.

### Stereotaxic surgery in mice

Mice (*Calca^Cre:GFP/+^*) were anesthetized with isoflurane (1-4%) and mounted in a stereotaxic frame (Kopf). Using standard surgical techniques, virus (AAV-DIO-GFP:TetTox or AAV-DIO-GFP) was injected bilaterally via a glass micropipette attached to a microinjector (Nanoject II) targeting the PBN (AP −4.8 mm; ML ±1.3 mm, DV −3.3 mm, relative to bregma).

### High-fat diet feeding

GFRAL^Cre-Dq^ mice were fed a high-fat diet (60% fat by kCal; research diets D12492) from weaning to 8 weeks of age prior to study. For GFRAL^CreERT-Dq^ studies, mice were fed high-fat diet from weaning, administered tamoxifen at 5 weeks of age and given an additional 5 weeks of HFD feeding prior to studies.

### Tamoxifen dosing

Animals were dosed at 8 weeks of age (except for the case of high fat diet fed animals, which were administered tamoxifen at 5 weeks of age) with 150 mg/kg tamoxifen dissolved in corn oil, delivered IP once per day for 5 days. Controls were vehicle treated. All experiments using *Gfral^CreERT^* were performed for a minimum of three weeks post-tamoxifen administration.

### Conditioned taste aversion

Mice were housed in custom cages with two ports on the front of the cage that accepted drinking bottles constructed of glass test tubes fitted with a rubber stopper and spout. On days 1-5 mice were injected with saline (0.01 ml/kg, sc) once daily. On the afternoon of day five the standard home cage water bottle was removed and replaced by custom drinking bottles filled with water. On day 8 water bottles were removed 1 hour before lights out. On day 9 a single pre-weighted bottle filled with 0.15% saccharin was offered 2 hours prior to lights out. At lights out on day 9, saccharin was removed and animals were injected with either vehicle or compound (GDF-15 (0.4 mg/kg, delivered sc; Novo Nordisk) or CNO 1 mg/kg ip; Tocris Bioscience) and water was returned. On day 10 water was removed from the cage 1 hour before lights out. On day 11, 1 hour before lights out two pre-weighed bottles were placed in the cage ports, one containing water and the other 0.15% saccharin. Bottles were weighed after 4 and 24 hours.

### Food-intake studies

Mice were singly housed a minimum of one week prior to studies. For acute studies, on the day of the experiment, food was removed at 4 PM and mice were randomized between CNO (1 mg/kg) or vehicle (sterile saline 0.9%) treatment. CNO or vehicle was administered via i.p. injection 30 minutes prior to the onset of dark and the food hopper was returned with a pre-weighed amount of food (generally ~10 g) at the onset of lights-out. Food was weighed 1,2, 4, 16 and 24 hours following the onset of the dark phase. Mice were given one week of rest, prior to the repetition of the experiment under inversed treatment conditions. In a subset of experiments, water intake was measured by the same experimental paradigm. Controls were either mice that did not express either Cre or hM3Dq. For chronic CNO dosing, singly housed mice were administered CNO once at 9 AM and once at 5 PM for up to 4 days. Food and body weight were monitored daily.

### CNO in drinking water

CNO was dissolved in water (supplemented with 1 % glucose) at a concentration of 2.5 mg/100mL H_2_O. Mice were singly housed a minimum of one week prior to experiments. Regularly supplied water was removed and replaced with water bottles containing CNO laced water as the only source of drinking water. Water, food consumption and mouse body weight were monitored daily. Water was replaced daily.

### Gastric Emptying

Mice were fasted four hours prior to CNO injections (1 mg/kg ip; Tocris Bioscience). Thirty minutes following CNO injection, mice were gavaged with 100 mg/kg acetaminophen dissolved in water and tail blood samples were collected for acetaminophen assay (Sekisui Diagnostics 506-30 Acetaminophen L3k Assay) 15 minutes following gavage.

### FOS studies

Liraglutide (400 μg/kg; delivered ip; Novo Nordisk), salmon calcitonin (400 μg/kg; delivered ip; Bachem), LiCl (84 mg/kg delivered ip; Sigma) and LPS (50 μg/kg delivered ip; Sigma) were injected into *Gfral^Cre^;Rosa26^eGFp-L10a/+^* mice two hours prior to sacrifice. Additionally, a separate cohort was fasted 16 hours (5 PM to 9 AM) and permitted two hours of food access prior to sacrifice, controls for refed mice were mice euthanized under fasting conditions. Mice were given GDF-15 (400 μg/kg; delivered sc; Novo Nordisk) and euthanized and perfused after 4 hours. Following fixation and staining, FOS+ cells were quantified in ImageJ within the AP and normalized to the number of GFP+ cells.

### In situ hybridizations

Mice were anesthetized with isoflurane and euthanized by decapitation. Brains were dissected, flash frozen in isopentane, chilled on dry ice, and stored at −80°C. Sections were sliced at 16 mm thickness using a cryostat (Leica) and every fourth section was thaw-mounted onto slides, allowed to dry for one hour at −20°C, and then further stored at −80°C. Slides were then processed for RNAScope ISH per the manufacturer’s protocol (Advanced Cell Diagnostics). The multiplex fluorescent assay (320850) was used to visualize *Gfral* (439141-C2) and *Cre* (312281-C3) probes using Amp 4 Alt-A. At each of 8 coronal planes for each mouse, 4 images comprising the entire NTS/AP complex were obtained with a QImaging Retiga 6000 monochrome camera attached to an Olympus BX53 fluorescent microscope under 20X objective. The four images were then stitched together using Photoshop (Adobe). CellProfiler was used to process all images identically to remove nonspecific background, outline specific cells using the DAPI nuclear signal and analyze presence or absence of signal for both probes. For each mouse, the total number of cells identified as positive for either or both probes were added from all 8 coronal planes (for NTS) or for the subset of the 6 planes where AP was present. Subsequently, the sums from the three mice were averaged for each region.

### Antibody staining

Mice were euthanized with isofluorane and then perfused by gravity flow with phosphate buffered saline for five minutes followed by an additional five minutes of 10% formalin fixation. Brains were then removed and post-fixed in 10% formalin for 24 hours at room temperature, before being moved to 30% sucrose for a minimum of 24 hours at room temperature. Brains were then sectioned 30-μm thick, free-floating sections and stained. Sections were treated sequentially with 1% hydrogen peroxide/0.5% sodium hydroxide, 0.3% glycine, 0.03% sodium dodecyl sulfate, and blocking solution (PBS with 0.1% triton, 3% normal donkey serum; Fisher Scientific). The sections were incubated overnight at room temperature in rabbit anti-FOS (FOS, #2250, Cell Signaling Technology, 1:1000; HA #C29F4, Cell Signaling Technology 1:1000; CGRP ab81887, Abcam, 1:2000). The following day, sections were washed and incubated with either biotinylated (1:200 followed by avidin-biotin complex (ABC) amplification and 3,3-diaminobenzidine (Thermo scientific) or fluorescent secondary antibody to visualize proteins. Immunofluorescent staining was performed using primary antibodies (GFP, GFP1020, Aves Laboratories; DSRed, 632392, Clontech) antibodies were reacted with species-specific Alexa Fluor-488, 568 conjugated secondary antibodies (Invitrogen, Thermo Fisher,1:300). Images were collected on an Olympus BX51 microscope. Images were pseudocolored and cell counts performed using ImageJ (NIH).

### Quantification and statistical analysis

All data are displayed as mean +/- SEM. Statistical analysis was performed in either Graphpad Prism 8 or R (specifically for Figure 2 O, P) using either t-tests, ANOVAs with Dunnet’s post-hoc test when appropriate or linear mixed model. P<0.05 was considered significant.

**Figure 1:**
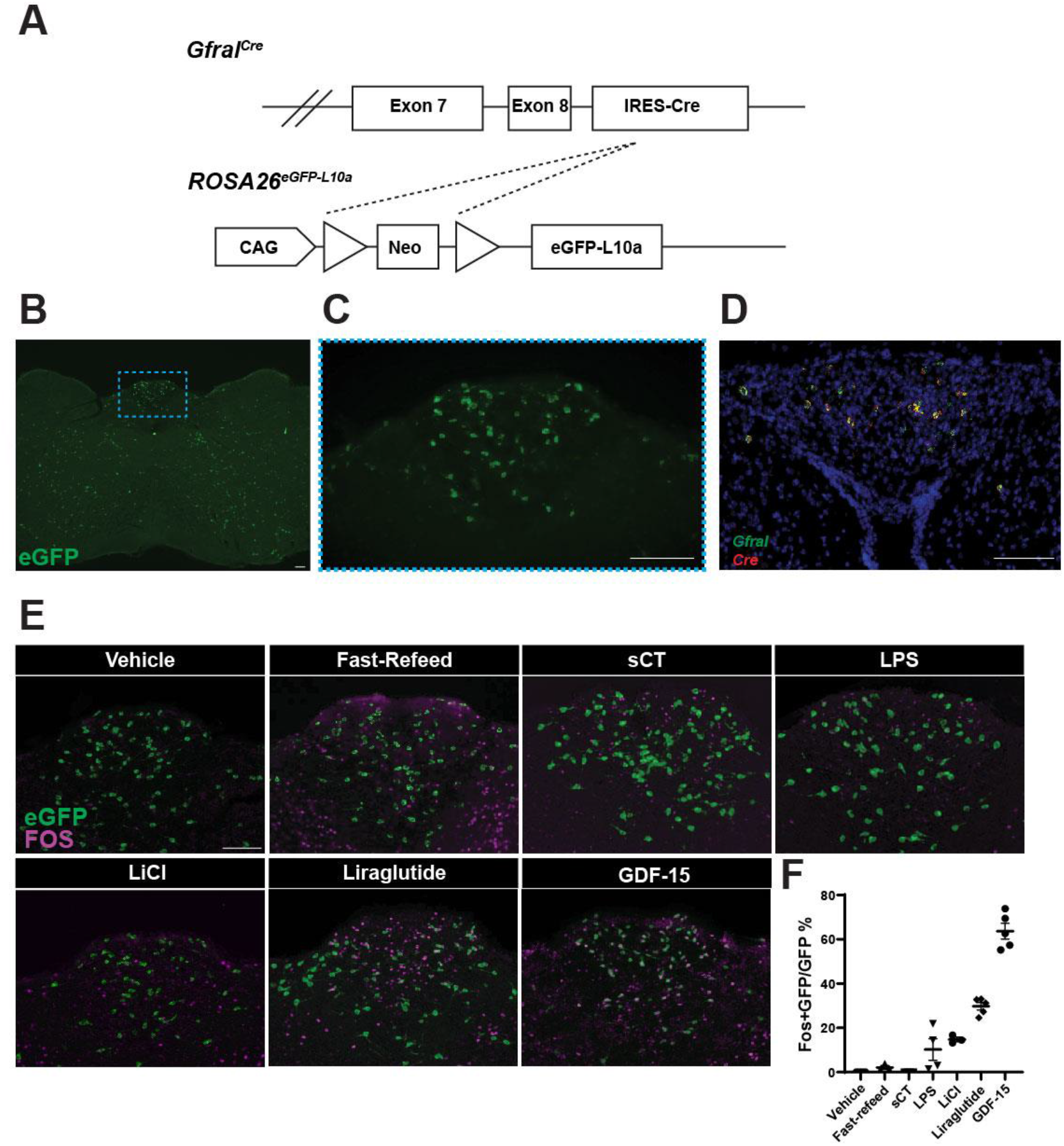
GDF-15 activates GFRAL neurons. (A) Schematic of the *Gfral^Cre^* -mediated excision of the Lox-Stop-Lox cassette from the *ROSA26^eGFP-L10a^* allele to generate GFRAL^eGFP^ mice. (B) Representative image of coronal hindbrain (Bregma −7.5) eGFP-IR (green) in GFRAL^eGFP^ mice. (C) High magnification of boxed AP region in (B). (D) Representative image of *in situ* hybridization for *Gfral* (Green) and *Cre* (Red) transcripts in AP/NTS of *Gfral^Cre^* mice. (E-F) Representative image and quantification of FOS-IR (Magenta) in GFRAL (eGFP+; Green) neurons from GFRAL^eGFP^ mice in response to Vehicle, Refeeding, sCT, LPS, LiCl, Liraglutide, and GDF-15 (n=3-6). Data are shown as mean ± SEM. Scale bar = 100 μm.

**Figure 2:**
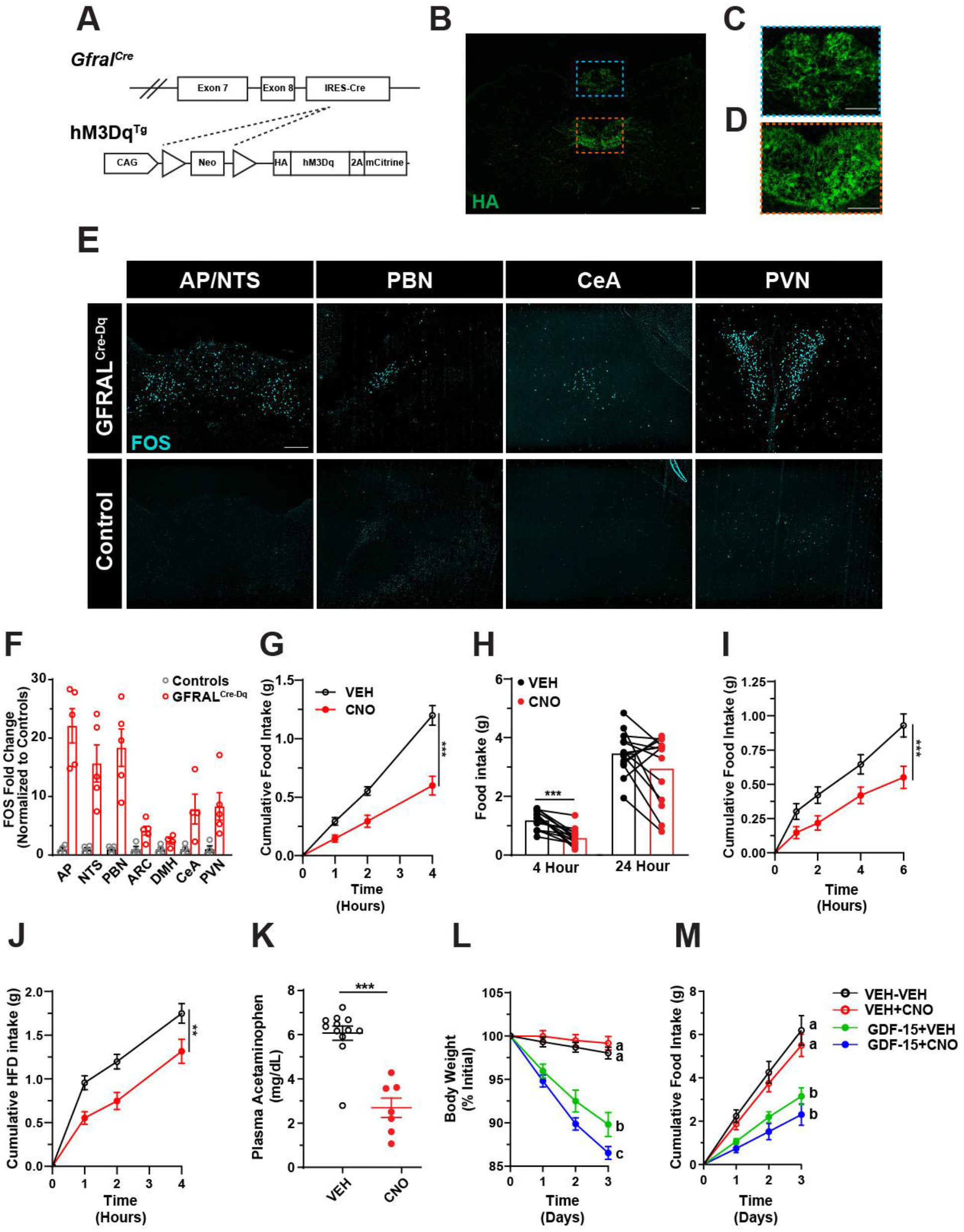
DREADD-mediated activation of *Gfral^cre^* neurons suppresses food intake and reduces body weight. (A) Schematic of *Gfral^Cre^*-mediated excision of the Lox-Stop-Lox cassette from the hM3Dq^Tg^ line to produce GFRAL^Cre-Dq^ mice. (B-C) Representative image of HA-IR (Green) in coronal hindbrain sections (Bregma −7.5) from GFRAL^Cre-Dq^ mice and higher magnification of AP (C, region in blue dashed box) and hypoglossal nucleus (D, region in orange dashed box). (E-F) Representative image (E) and quantification (F) of FOS-IR (Cyan) in the AP/NTS, PBN, CeA and PVN from CNO-treated (1mg/kg; ip) GFRAL^Cre-Dq^ and control animals (n=4). (G) Food intake over the first 4 hours of the dark phase in GFRAL^Cre-Dq^ mice following either vehicle (VEH; black lines) or CNO (red lines) injection 30 minutes prior to the onset of the dark cycle (n=14). (H) Paired analysis of 4- and 24-hour food intake in GFRAL^Cre-Dq^ animals administered CNO or VEH. (I) Light phase food intake levels following a 10-hour overnight fast in GFRAL^Cre-Dq^ mice administered vehicle or CNO (n=18). (J) Dark phase food intake of a palatable diet (60% fat) in lean GFRAL^Cre-Dq^ animals injected with either vehicle or CNO (n=14). (K) Plasma acetaminophen concentrations 30 minutes after oral gavage in controls and GFRAL^Cre-Dq^ animals administered CNO (n=7-12). (L-M) Body weight and food intake of DIO GFRAL^Cre-Dq^ animals injected twice daily with either vehicle or CNO in the presence or absence of a single daily injection of GDF-15 (n=11). Data are shown as mean ± SEM. Two-way ANOVA (G,I,J), paired t-test (H), unpaired t-test (K), or linear mixed model (L,M) was performed, **p < 0.01, ***p < 0.001. Scale bar= 100 μm.

### Data and Code Availability

This study did not generate any unique datasets or code.

## Results

While recent studies have elucidated some aspects of the biology of GDF-15 and its receptor, GFRAL, little is known about the regulation, function, and downstream mediators of GFRAL-expressing neurons. To study GFRAL neurons independently of GDF-15 biology, we developed a *Gfral^Cre^* knock-in mouse allele. Breeding *Gfral^Cre^* onto the Cre-inducible *ROSA26^eGFP-L10a^* reporter background (GFRAL^eGFP^ mice) revealed the expected presence of many eGFP-expressing neurons within the AP, as well as scattered eGFP-containing cells elsewhere within the hindbrain (Figure 1B, C). *In situ* hybridization revealed the restriction of *Gfral* expression to the AP/NTS (in agreement with previous observations (Emmerson et al., 2017; Hsu et al., 2017; Mullican et al., 2017; Yang et al., 2017)), where it colocalized with *Cre* mRNA (Figure 1D). The presence of eGFP reporter expression outside of the AP/NTS presumably indicates some transient developmental expression of *Gfral^Cre^* in these areas. Consistent with a crucial role for AP GFRAL neurons in the control of food intake, direct GDF-15 administration to rat AP (but not the NTS) produced a strong anorexic response (Supplemental Figure 1A, B).

We first sought to define the types of stimuli that activate GFRAL neurons. Unlike other hindbrain cells known to modulate food intake (Cheng et al., 2020b, 2020a; Roman et al., 2016) refeeding following an overnight fast failed to promote the accumulation of FOS-immunoreactivity (-IR) in eGFP-labelled AP GFRAL neurons in GFRAL^eGFP^ mice; neither did treatment with the appetite-suppressing calcitonin receptor (CALCR) agonist, salmon calcitonin (sCT). In contrast, lipopolysaccharide (LPS), liraglutide (a glucagon-like peptide-1 receptor (GLP-1R) agonist), LiCl (which causes GI distress), and GDF-15 promoted FOS-IR in many AP GFRAL cells (Figure 1E, F). These data suggest that GFRAL neurons do not respond to meal-related signals and that GFRAL cells are distinct from CALCR neurons which respond to sCT and meal-related cues (Cheng et al., 2020). Instead, GFRAL neurons respond to signals associated with pathophysiology, including GDF-15, GI distress, bacterial infection, and GLP-1 system stimulation (systemic inflammation, as during a variety of infections, activates the endogenous GLP-1 system (Lebrun et al., 2017)).

Because manipulation of neurons in the mouse AP by the stereotaxic injection of viral vectors has proved difficult, we took advantage of the restricted *Gfral* expression pattern and utilized genetic systems to study AP GFRAL neurons. To study the functional capacity of AP GFRAL neurons, we crossed *Gfral^Cre^* mice onto the Cre-inducible hM3Dq^Tg^ background (Zhu et al., 2016) to express the activating (hM3Dq) designer receptor exclusively activated by designer drugs (DREADD) in GFRAL neurons (GFRAL^Cre-Dq^ mice), permitting the activation of these cells by CNO administration (Figure 2A). While we observed strong DREADD expression within the AP of GFRAL^Cre-Dq^ mice, we also noted DREADD expression outside of the AP, most densely in the hypoglossal and facial nuclei (Figure 2B-D; Supplemental Figure 2A). CNO injection in GFRAL^Cre-Dq^ mice increased FOS-IR within the AP, consistent with the expected activation of hM3Dq-expressing GFRAL neurons by CNO. Notably, a number of other regions also displayed increased FOS-IR, including the NTS, PBN, CeA and PVN (Figure 2E, F). These regions likely lie downstream of GFRAL neurons, as GDF1-15 also promotes FOS-IR in these areas (Johnen et al., 2007).

To determine the ability of GFRAL neuron activation to modulate feeding behavior, we administered CNO 30 minutes prior to the onset of the dark cycle and monitored food intake. In GFRAL^Cre-Dq^ mice, CNO treatment dramatically suppressed food intake compared to saline injection, although this effect was attenuated by 24 hours (Figure 2G,H). Importantly, CNO did not alter food intake in control animals (lacking either Cre or DREADD expression; Supplemental Figure 2B-D). We also examined the ability of GFRAL neuron activation to suppress food intake in the face of an elevated drive to feed. Following a 10-hour fast or acute exposure to palatable food (60% fat diet), activation of GFRAL neurons by CNO treatment decreased food intake by approximately 50% (Figure 2I, J). We also found that CNO treatment of GFRAL^Cre-Dq^ mice decreased gastric emptying (Figure 2K).

To determine whether chronic activation of GFRAL neurons could suppress food intake and decrease body weight over longer periods of time, we treated GFRAL^Cre-Dq^ mice with CNO over three days (with or without an additional daily injection of GDF-15). This CNO-dosing scheme mediated the continued suppression of food intake and body weight over the three-day duration of the experiment. Interestingly, exogenous GDF-15 amplified CNO-mediated weight loss (Figure 2L,M), suggesting that GDF-15 does more to the GFRAL neuron than increasing its activity. Altogether, these data reveal that activating GFRAL neurons reduces food intake and body weight more effectively than GDF-15, although GDF-15 amplifies the effects of GFRAL neuron activation, presumably by altering other aspects of these neurons. GFRAL neurons also decrease gastric emptying, suggesting that they limit nutrient absorption as well as food intake.

In a separate experiment initially designed as a chronic CNO-dosing paradigm, we provided mice with CNO-laced glucose-containing water as their only source of drinking water. To our surprise, GFRAL^Cre-Dq^ mice suppressed their consumption of CNO- and glucose-containing water compared to control mice (which received the same CNO- and glucose-containing drinking water) to a greater extent than they decreased food intake. CNO also reduced body weight in GFRAL^Cre-Dq^ mice (Figure 3A-C). Injection of CNO (ip) did not decrease water consumption, however (Supplemental Figure 4D-F). Thus, GFRAL neuron activation does not simply suppress thirst. These experiments suggested that the activation of GFRAL neurons is aversive.

**Figure 3:**
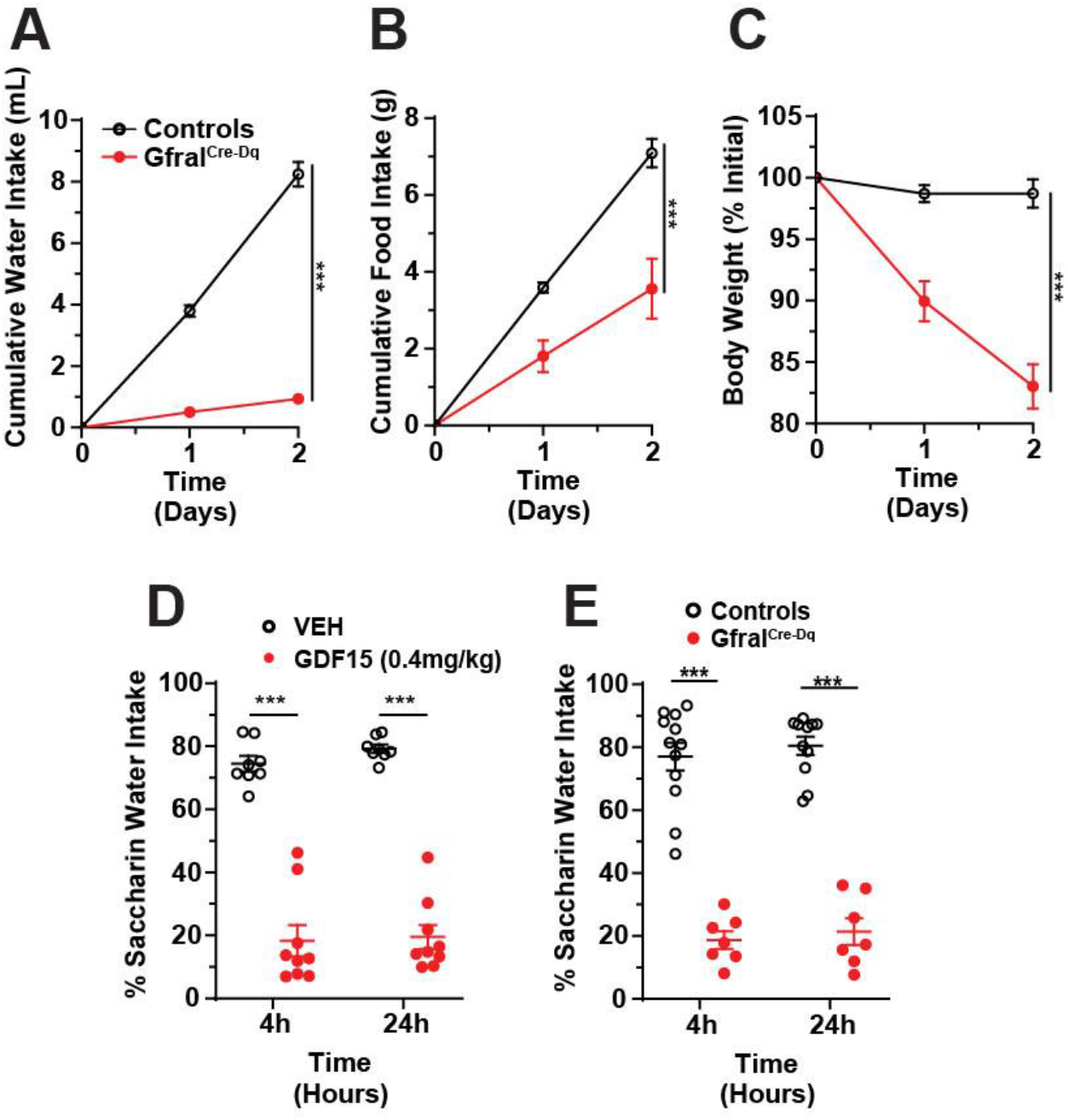
Activation of GFRAL neurons, like GDF-15, promotes aversive responses. (A-C) Water intake (A), food intake (B), and body weight (C) in control and GFRAL^Cre-Dq^ mice over two days during which time the only source of drinking water contained CNO (0.025 mg/mL; n=6). (D) Conditioned taste aversion (CTA) assay in wild-type mice injected with either vehicle (VEH) or GDF-15 during conditioning (n=8-9). (E) CTA in GFRAL^Cre-Dq^ mice and Controls injected with CNO in conditioning phase (n= 7-12). Data are shown as mean ± SEM. Two-way ANOVA (A,B,C,D,E) was performed, ***p < 0.001.

To test whether GDF-15 signaling can provoke a conditioned taste aversion (CTA), we gave mice saccharin in addition to GDF-15 or vehicle and then tested them the next day for their preference of saccharin or water. We found GDF-15 produced a strong CTA (Figure 3D), consistent with the findings of others (Borner et al., 2020; Patel et al., 2019). Furthermore, artificial activation of GFRAL neurons by treating GFRAL^Cre-Dq^ mice with CNO also produced a strong CTA to saccharin (Figure 3E). These studies show that GDF-15 and artificial activation of GFRAL neurons mimics gastrointestinal malaise in the CTA assay.

While these studies suggest the ability of GFRAL neurons to suppress food intake and promote aversive responses independently of GDF-15 signaling, the expression of the hM3Dq in cells outside the AP in GFRAL^Cre-Dq^ mice complicates the interpretation of these studies (although neither the hypoglossal or facial nucleus have been implicated in either feeding or aversive behaviors). Because we observed *Gfral^Cre^*-mediated recombination in regions inconsistent with adult *Gfral* expression patterns in both Cre-dependent GFP and hM3Dq alleles, we speculated that the transient developmental expression of *Gfral* (and *Gfral^Cre^*) might mediate recombination outside of the AP. To overcome this concern, we generated a second *Gfral* knock-in allele to express tamoxifen-inducible CreERT (*Gfral^CreERT^*; Figure 4A) to permit the manipulation of GFRAL neurons in adulthood.

**Figure 4:**
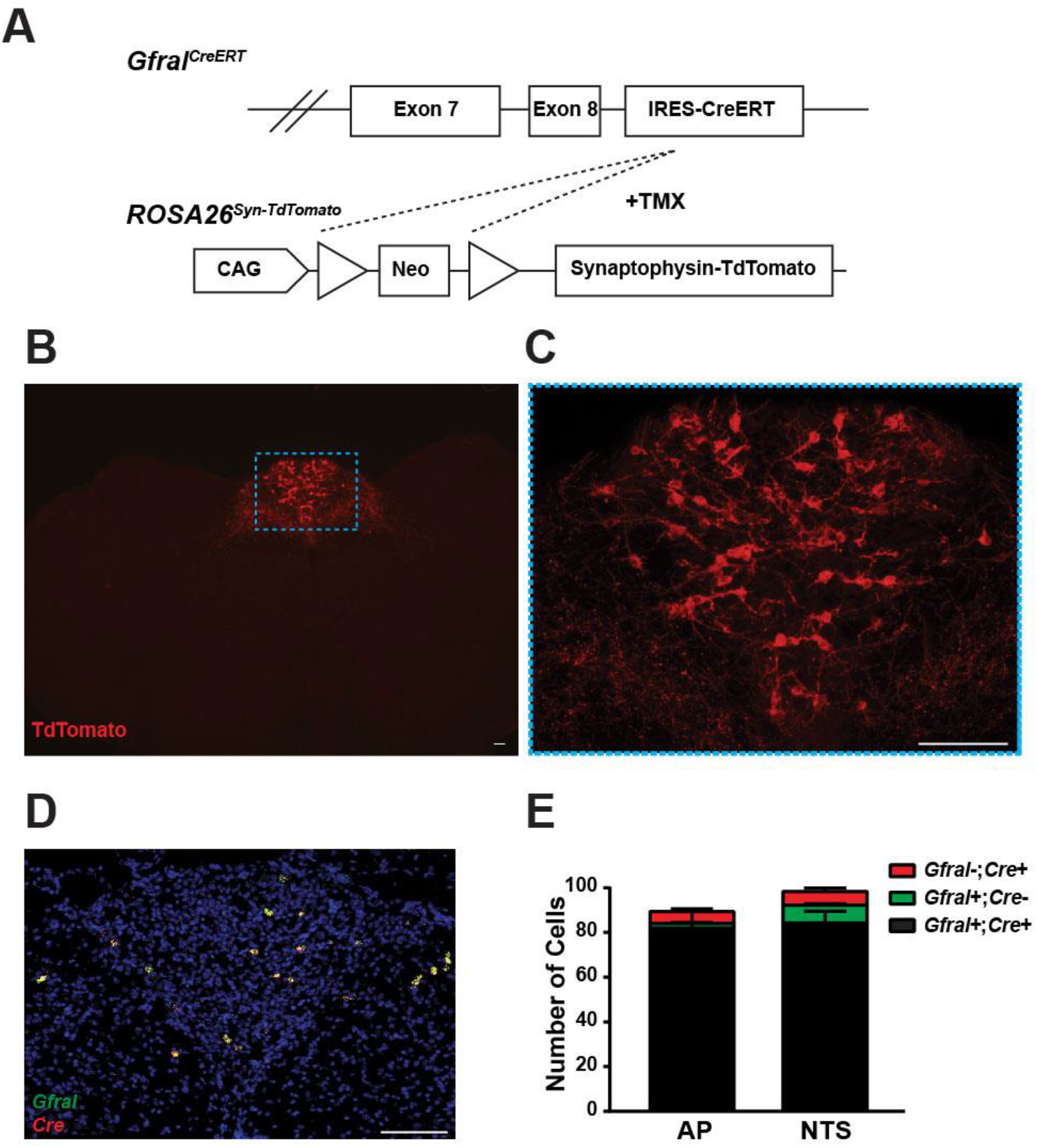
Generation of a tamoxifen-inducible *Gfral^CreERT^* allele. (A) Schematic of *Gfral^CreERT^* – and tamoxifen (TMX)-mediated recombination of the Lox-Stop-Lox cassette from the *ROSA26^Syn-TdTomato^* line to generate GFRAL^CreERT-TdTomato^ mice. (B-C) Representative image of TdTomato-IR (Red) in coronal hindbrain (Bregma −7.5) sections from GFRAL^CreERT-TdTomato^ mice three weeks post TMX treatment. (C) Shows high magnification of the boxed AP region in (B). (D-E) Representative image and quantification of *in situ* hybridizations for *Cre* (Red) and *Gfral* (Green) transcripts in the AP/NTS of *Gfral^CreERT/CreERT^* mice (n=3). Data are shown as mean ± SEM. Scale bar = 100 μm.

We crossed *Gfral^CreERT^* onto the Cre-inducible *ROSA26^Syn-TdTomato^* reporter background (Figure 4A) and treated the resultant animals with tamoxifen (TMX), which promoted somatic TdTomato expression exclusively within the AP (Figure 4B, C), albeit in fewer AP cells than observed with the standard *Gfral^Cre^* mouse. *In situ* hybridization confirmed a high degree of co-localization between *Cre* and *Gfral* transcripts in this model (Figure 4D, E).

We crossed *Gfral^CreERT^* on to the Cre-inducible hM3Dq^Tg^ background and treated the resultant mice with TMX (GFRAL^CreERT-Dq^ mice) to permit activation of AP GFRAL neurons specifically (Figure 5A). As with the TdTomato reporter, the *Gfral^CreERT^* allele promoted hM3Dq expression only in the AP of GFRAL^CreERT-Dq^ mice, although *Gfral^CreERT^* mediated hM3Dq expression in fewer AP cells than *Gfral^Cre^* (despite utilizing *Gfral^CreERT^* in the homozygous state) (Figure 5B, C; Supplemental Figure 3A, B). As for GFRAL^Cre-Dq^ mice, CNO administration in GFRAL^CreERT-Dq^ mice promoted FOS-IR in the AP, NTS and exterior lateral PBN; while FOS-IR also tended to increase in the CeA and PVN, the effect was not statistically significant in these regions (perhaps due to the decreased number of GFRAL neurons targeted in GFRAL^CreERT-Dq^ compared to GFRAL^Cre-Dq^ mice) (Figure 5D, E).

**Figure 5:**
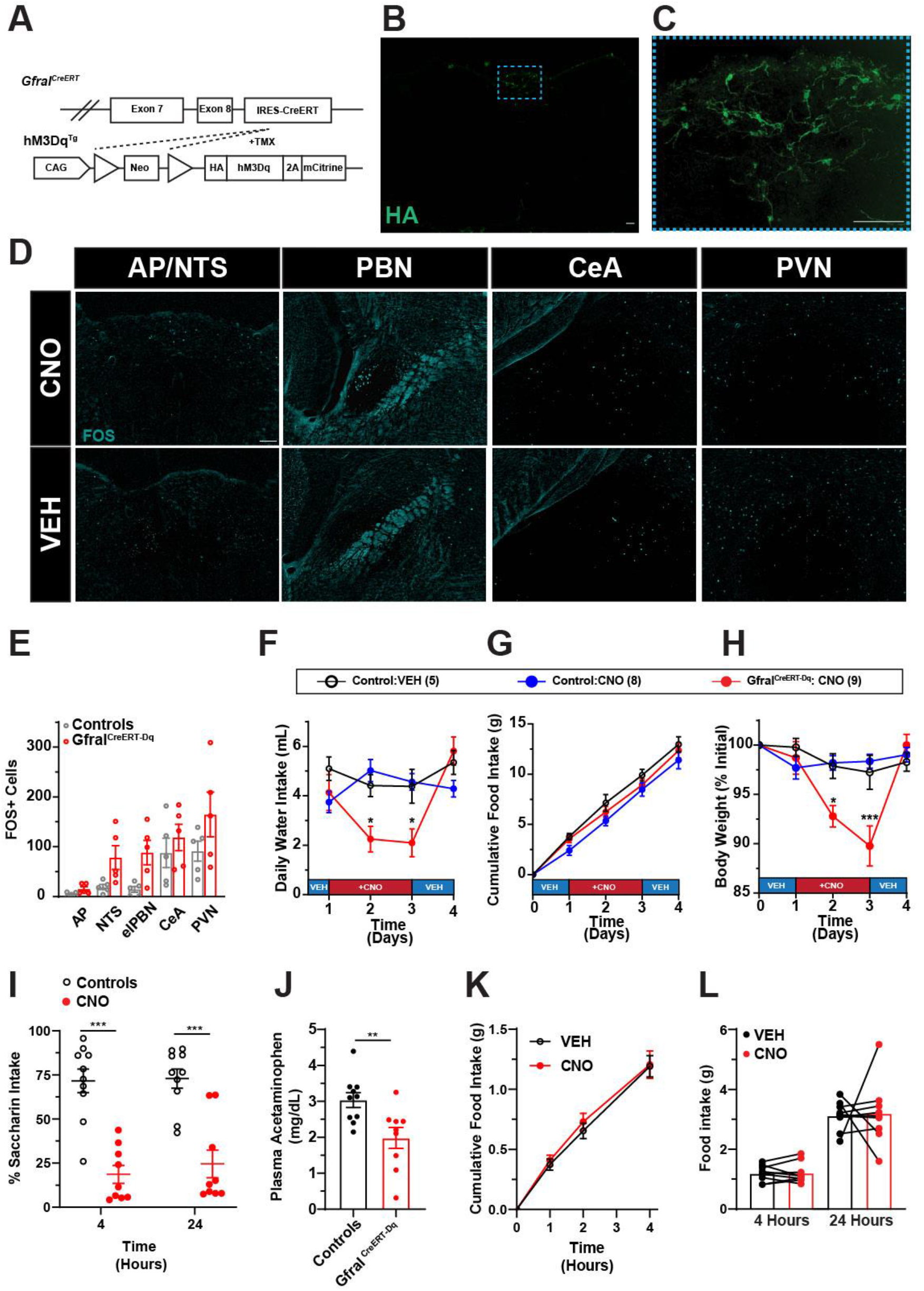
Activation of *Gfral^CreERT^* neurons is aversive but does not decrease food intake. (A) Schematic of *Gfral^CreERT^*- and tamoxifen (TMX)-mediated recombination of the Lox-Stop-Lox cassette from the hM3Dq^Tg^ line to generate GFRAL^CreERT-Dq^ mice. (B-C) Representative image of HA-IR (Green) in coronal hindbrain sections (Bregma −7.5). (C) Shows higher magnification of the AP region boxed in (B). (D-E) Representative image (D) and quantification (E) of FOS-IR (Cyan) in the AP/NTS, PBN, CeA and PVN two hours following CNO or vehicle (VEH) administration in GFRAL^CreERT-Dq^ animals (n=5). (F-H) Daily water intake (F), Cumulative food intake (G), and body weight (H) GFRAL^CreERT-Dq^ animals exposed to control water (days 0-1 & 3-4) and CNO-laced drinking water (days 1-3). Control animals were divided between those that received glucose-containing water alone (VEH) on days 1-3 and those that received CNO-laced, glucose-containing water on days 1-3 (CNO) (n values in parenthesis in figure legend). (I) Conditioned taste aversion in GFRAL^CreERT-Dq^ mice and controls injected with CNO during conditioning (n= 9-10). (J) Plasma acetaminophen levels 30 minutes following oral gavage of 100mg/kg acetaminophen in control or GFRAL^CreERT-Dq^ mice administered CNO. (K) Food intake over the first four hours of the dark phase in GFRAL^CreERT-Dq^ mice following either vehicle (VEH) or CNO injection 30 minutes prior to the onset of the dark cycle (n=9). (L) Paired analysis of 4- and 24-hour food intake in GFRAL^CreERT-Dq^ animals administered CNO or VEH. Data are shown as mean ± SEM. Two-way ANOVA (F,G,H,I,K), unpaired t-test (J) or paired t-test (L) were performed, *>0.05, **p < 0.01, ***p < 0.001. Scale bar = 100 μm.

Despite having fewer reporter-labelled cells and less robust FOS responses to CNO compared to GFRAL^Cre-Dq^ mice, GFRAL^CreERT-Dq^ mice responded to CNO in their drinking water by reducing water intake and body weight over the two-day treatment, albeit without altering food intake (Figure 5F-H). We observed similar results in multiple cohorts of GFRAL^CreERT-Dq^ mice, including those that were only heterozygous for *Gfral^CreERT^* (Supplemental Figure 4M-O). Additionally, CNO provoked a CTA to saccharin-flavored water and slowed gastric emptying in GFRAL^CreERT-Dq^ mice (Figure 5I, J).

Importantly, however, while the aversive and gastric emptying effects of CNO in GFRAL^CreERT-Dq^ mice resembled those of GFRAL^Cre-Dq^ mice, CNO failed to suppress food intake in GFRAL^CreERT-Dq^ mice (Figure 5K-L). CNO also failed to suppress food intake in a separate cohort of GFRAL^CreERT-Dq^ mice fed a high-fat diet to induce diet-induced obesity (Supplemental Figure 4J). Furthermore, twice-daily CNO injections for four days failed to suppress body weight or food intake in GFRAL^CreERT-Dq^ mice (Figure Supplemental Figure 4K, L). These results suggest that the suppression of food intake requires the activation of greater numbers of GFRAL neurons than is required to produce a CTA or alter gastric emptying. Indeed, the number of HA-tagged DREADD expressing GFRAL neurons detected in the GFRAL^CreERT-Dq^ mice correlated inversely with CNO-laced water intake, but not with dark-phase food intake, during CNO treatment (Supplemental Figure 4Q, R)

To understand the downstream circuits by which GFRAL neurons act, we crossed *Gfral^CreERT^* onto the Cre-inducible *ROSA26^Syn-TdTomato^* background to express the synaptophysin-TdTomato fusion protein in GFRAL neurons (Figure 6A), allowing us to visualize their projections. Following TMX administration, we observed somatic reporter expression only within the AP, consistent with the specificity of synaptophysin-TdTomato expression in AP GFRAL neurons in this model (Figure 6B). Examining the entire brain for TdTomato revealed projections only in the NTS and the external lateral PBN. The PBN terminals lie in close proximity to CGRP^PBN^ neurons (Figure 6C).

**Figure 6:**
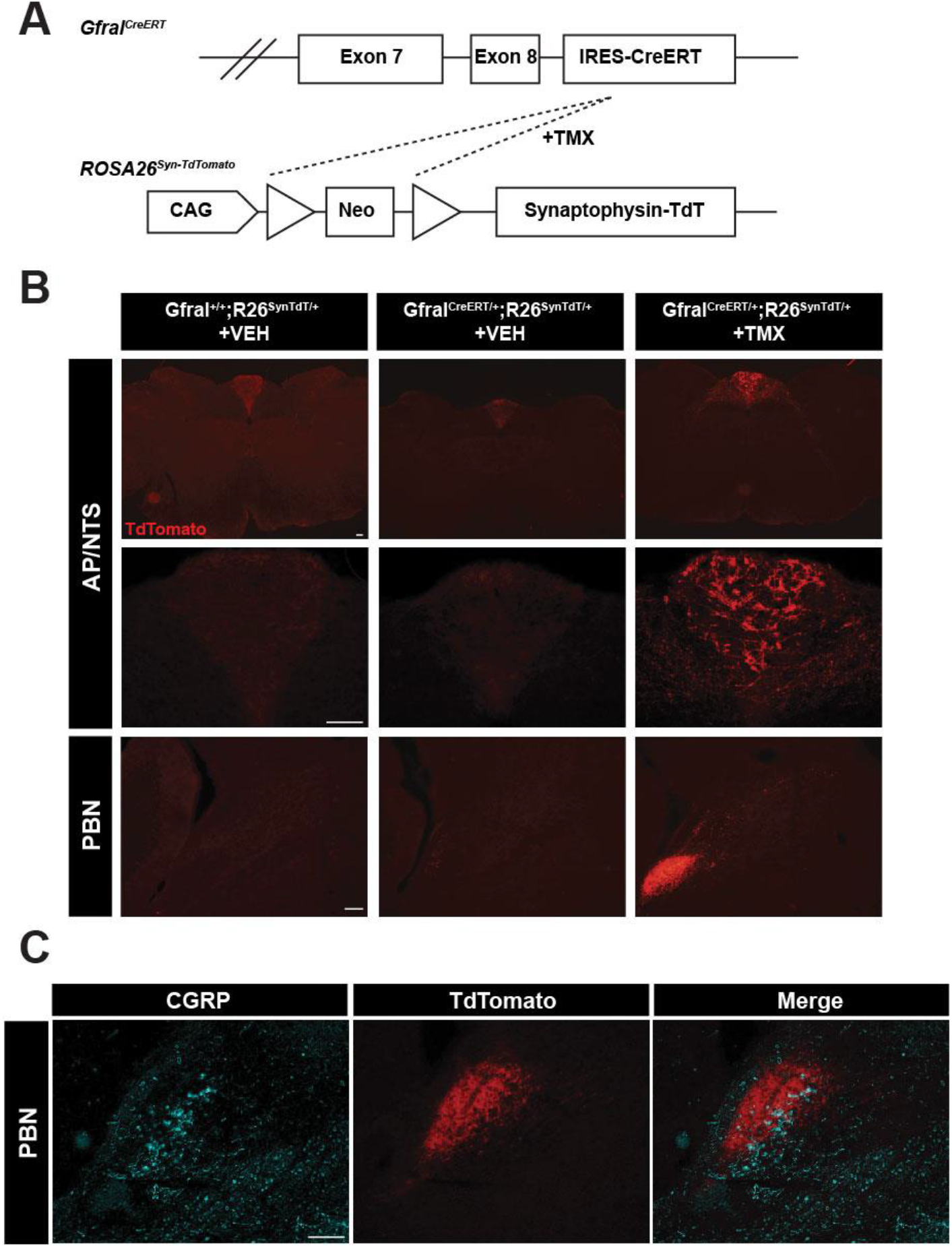
GFRAL neurons project to CGRP^PBN^ neurons. (A) Schematic of the *Gfral^CreERT^*- and tamoxifen (TMX)-mediated recombination of the Lox-Stop-Lox cassette from *ROSA26^SynTdT^* to generate GFRAL^CreERT-SynTdT^ mice. (B) Representative image of TdTomato-IR (Red) in AP/NTS and PBN of *Gfral^+/+^* mice, GFRAL^CreERT-SynTdT^ mice and GFRAL^CreERT-SynTdT^ mice administered TMX or vehicle (VEH), as indicated. (C) Representative image of TdTomato- (Red) and CGRP-IR (Cyan) within the PBN from TMX-treated GFRAL^CreERT-SynTdT^ mice. Scale bar = 100 μm.

As CGRP^PBN^ neurons have previously been described to mediate aversive and anorectic outputs (Carter et al., 2013, 2015), we hypothesized that this PBN cell population might be required for GDF-15 action. Indeed, GDF-15 promoted FOS-IR in ~40% of CGRP neurons in *Calca^Cre:GFP^* mice (Figure 7A-B). To test the requirement for CGRP^PBN^ neurons in GDF-15 action, we injected the Cre-dependent AAV DIO-GFPTetTox into the PBN of *Calca^Cre:GFP^* mice (CGRP^PBN-TetTox^ mice) to express the light chain of tetanus toxin in CGRP^PBN^ cells, silencing them (Figure 7C). We found that GDF-15 failed to promote a CTA to saccharine-laced water in CGRP^PBN-TetTox^ mice and that CGRP^PBN-TetTox^ mice exhibited an attenuated anorectic response to GDF-15 (Figure 7D, E) indicating CGRP^PBN^ neurons mediate the aversive and anorectic responses to GDF-15.

**Figure 7:**
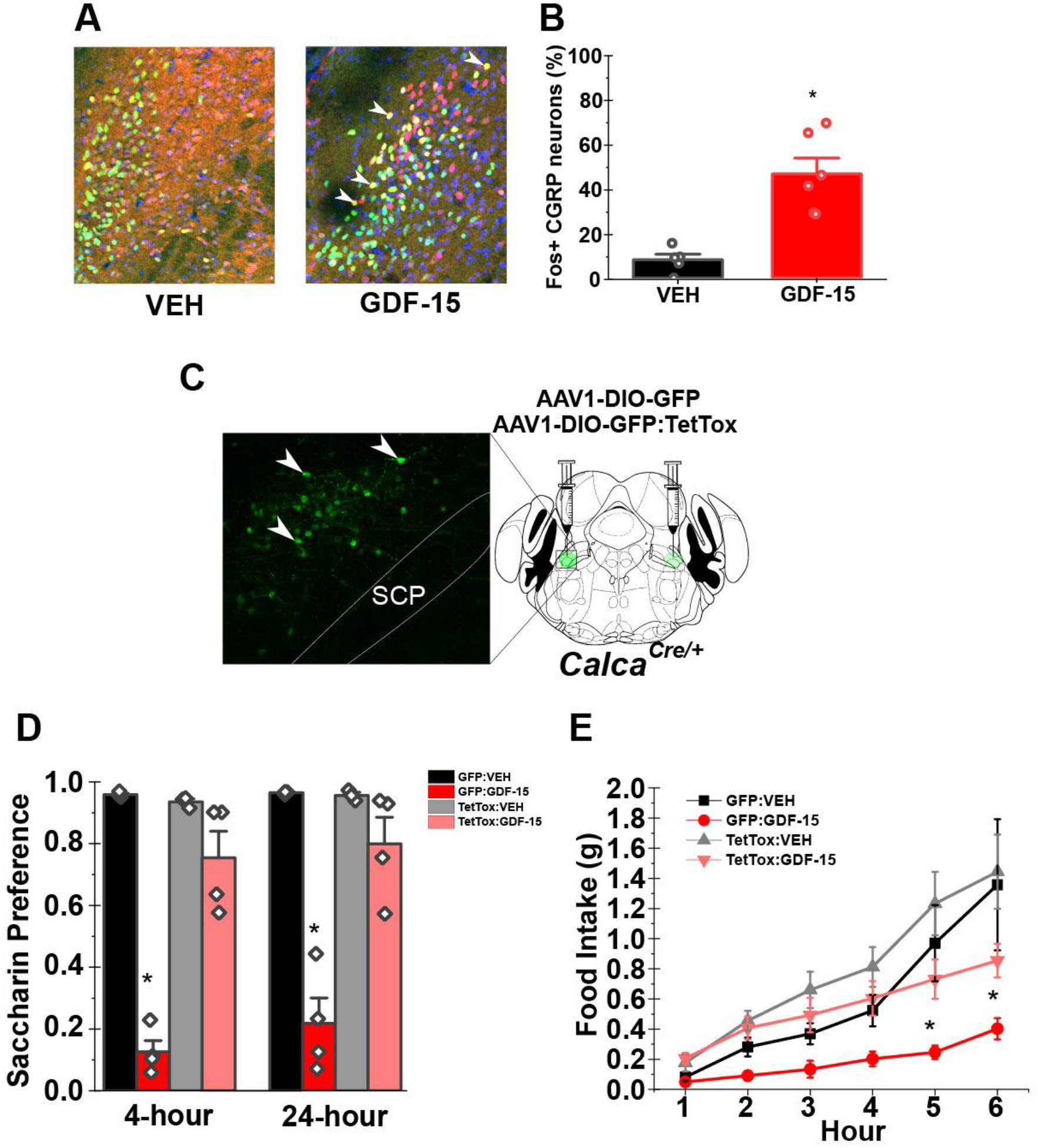
GDF-15-mediated feeding suppression and aversion require signaling by CGRP^PBN^ neurons. (A-B) Representative images (A) and quantification (B) of PBN FOS-IR four hours following vehicle (VEH) or GDF-15 (400 μg/kg) administration (DNA:blue, CGRP:green, FOS:Red; n=5). (C) Representative image of GFP-IR (Green) from the PBN of *Calca^Cre:GFP/+^* animals injected with AAV-DIO-GFP:TetTox and schematic of bilateral delivery methodology to the elPBN of *Calca^Cre:GFP/+^* animals. (D) CTA in AAV-DIO-GFP or AAV-DIO-GFP:TetTox injected *Calca^Cre:GFP/+^* mice administered with either vehicle or GDF-15 during conditioning (n=4). (E) Dark phase food intake of in AAV-DIO-GFP or AAV-DIO-GFP:TetTox injected *Calca^Cre:GFP^* mice injected with either vehicle or GDF-15 one hour prior to the onset of the dark cycle (n=5). SCP= superior cerebellar peduncle. Data are shown as mean ± SEM and analyzed by Two-way ANOVA. *p < 0.05

## Discussion

The important function and therapeutic potential of GDF-15, the hindbrain-restricted expression of GFRAL (Emmerson et al., 2017; Mullican et al., 2017; Yang et al., 2017), and the dearth of information about roles for AP neural populations in the control of food intake and body weight prompted us to study GFRAL neurons. In addition to using genetic approaches to confirm that GFRAL neurons reside predominantly in the AP, we demonstrate that AP GFRAL neurons mediate the anorectic effects of GDF-15. We also found that signals associated with systemic infection or illness stimulate GFRAL cells, while meal-related signals do not activate GFRAL neurons. Furthermore, GFRAL neurons inhibit gastric emptying, suppress food intake, and promote aversion, suggesting that GFRAL neurons attenuate the uptake of nutrients during pathophysiologic states. GFRAL neurons project most strongly to the PBN, where they innervate the CGRP^PBN^ cells that transmit negative-valence signals in response to a variety of pathophysiologic stimuli (Campos et al., 2017; Carter et al., 2013), including the aversive and anorectic responses to GDF-15. Overall, our findings suggest that GFRAL neurons detect pathophysiologic signals for relay to CGRP^PBN^ cells as part of the system that promotes aversion and attenuates nutrient uptake.

GFRAL neurons project to the PBN but not to hypothalamic sites that are also critical for homeostatic control of food intake and body weight. Indeed, GDF-15 levels are not regulated in a manner consistent with an endogenous regulator of body weight (Patel et al., 2019) and the lack of GDF-15 or GFRAL minimally, if at all, impacts body weight under normal conditions (Emmerson et al., 2017; Hsu et al., 2017; Mullican et al., 2017; Tran et al., 2018; Tsai et al., 2013; Yang et al., 2017). Similarly, CGRP^PBN^ neurons contribute to the decreased feeding and body weight associated with pathophysiologic states such as cancer, chemotherapy, or infection (Palmiter, 2018), but they are not essential for normal body weight regulation (Carter et al., 2013). Thus, GFRAL neurons appear to function primarily to regulate the control of feeding and GI function under pathophysiologic conditions.

Interestingly, GFRAL neurons decrease gastric emptying in a manner similar to other populations of hindbrain neurons that control feeding (Travagli et al., 2006). The projections from GFRAL neurons into the NTS may modulate the activity of neurons in the dorsal motor nucleus of the vagus to decrease gastric tone. The mechanism notwithstanding, this effect suggests that GFRAL neurons slow nutrient absorption, as well as intake, to decrease overall nutrient uptake by the body.

Acutely activating *Gfral^Cre^* neurons potently reduces food intake in a variety of feeding conditions (as well as promoting a CTA and decreasing gastric emptying), and chronic activation of *Gfral^Cre^* neurons results in substantial reductions in food intake and body weight. Despite being specific for GFRAL neurons within the AP and NTS, *Gfral^Cre^* mediates recombination and expression of hM3Dq^Tg^ in additional populations of neurons, including in the hypoglossal and facial nuclei. While little evidence links these extra-AP/NTS nuclei to the regulation of food intake, we repeated these functional studies with the TMX-inducible *Gfral^CreERT^* allele. While TMX treatment of *Gfral^CreERT^*-expressing mice yielded a more specific hM3Dq expression pattern (limited to the AP and a small number of cells in NTS), it did not promote recombination in all (or even the majority of) AP GFRAL cells. Less robust recombination mediated by a TMX-sensitive CreERT has been observed in other systems, as well (Thorens et al., 2015).

Interestingly, activation of this smaller number of AP/NTS cells sufficed to promote a robust CTA and to slow gastric emptying, but not to reduce food intake (see Figure 5). While this could reflect some unexpected contribution of the facial or hypoglossal nucleus to food intake, it is more likely that the failure of GFRAL neuron activation to suppress food intake in the conditional model reflects the failure to activate most of the AP GFRAL neurons in this model. Indeed, the difference in phenotypes between the *Gfral^Cre^* and *Gfral^CreERT^* models aligns with the dose-dependent behavioral responses to GDF-15 in rats, wherein low doses of GDF-15 induce pica behavior while considerably higher concentrations are required to suppress food intake (Borner et al., 2020). These data indicate that it takes considerably less activation of GFRAL neurons to produce a substantial aversive response than to produce anorexia, although we cannot rule out the possibility that *Gfral^CreERT^* mediates recombination in a specialized subset of GFRAL neurons.

GDF-15 promotes a sense of visceral malaise as assessed by CTA and other measures (Borner et al., 2020; Patel et al., 2019)(and our data). Consistent with this effect of GDF-15, acute activation of GFRAL neurons also promotes a strong CTA. This aversive effect presents a potential challenge to the use of GDF-15 or its analogs for the pharmacological treatment of obesity, since GDF-15 would be much less attractive to use if its weight loss effects result from a feeling of visceral illness. This represents the challenge of employing a target (such as the GDF-15/Gfral system) that is not part of the normal regulation of body weight, but rather modulates food intake under pathophysiological conditions. There exist other examples of pharmacological agents (such as long-acting GLP-1R agonists) that produce potent CTAs in rodents (Adams et al., 2018; Kanoski et al., 2012) and cause initial visceral illness in humans (Lean et al., 2014). However, most patients who continue on these drugs become tolerant to the nausea-inducing effects but continue to lose weight (Garber, 2011). Hence, the effects of GDF-15 on nausea may be separable from its effects on food intake and weight.

Indeed, several conditions that result in chronically elevated GDF-15 levels do not result in persistent nausea. Metformin results in increased GDF-15 levels and the increased GDF-15 plays a major role in the ability of metformin to cause weight loss (Coll et al., 2019), although most metformin-treated patients do not report persistent nausea. GDF-15 levels are also elevated in the second trimester of pregnancy (Petry et al., 2018). Nevertheless, the second trimester is actually associated with reduced visceral illness and emesis as compared to the first trimester for the majority of women (Eliakim et al., 2000).

In summary, our findings suggest that AP GFRAL neurons detect pathophysiologic signals for relay to CGRP^PBN^ cells as part of the system that promotes aversion and attenuates nutrient uptake during pathophysiologic states, rather than contributing to homeostatic energy balance. In the future, it will be particularly important to understand how GDF-15 may provoke activity-independent changes in GFRAL neurons and to define the roles for GFRAL neurons in mediating the aversive and food intake effects by non-GDF-15-related signals.

## Author Contributions

PVS, HFS, DG, AP, JM, JA, JW performed experiments and analyzed data. DPO provided crucial reagents. PVS, HFS, MGM, RDP,SBJ, RJS designed experiments and wrote and edited the manuscript. All authors reviewed and edited the manuscript. RJS is the guarantor of the manuscript.

## Declaration

SBJ is an employee of Novo Nordisk. DPO, MGM, and RJS receive research support from Novo Nordisk. RJS and MGM receive research support from AstraZeneca. RJS receives research support from Pfizer, Kintai and Ionis. RJS also serves as a paid consultant for Novo Nordisk, Kintai, Ionis and Scohia. RJS has equity positions in Zafgen and ReDesign Health. All other authors report no conflicts of interest.

## Acknowledgments

We acknowledge the technical assistance of the University of Michigan Transgenic Core. PVS was supported by the American Diabetes Association Grant 1 −19-PDF-099, JW was supported by China Scholarship Council. JMA was supported by T32-DK-101357. (This research was also supported in part by NIH grants (R01-DA-24908 to RDP, P30-DK089503, P01-DK117821, R01-DK119188 to RJS and P30-DK020572 to MGM).

