## Supplemental Figures 1-4 for "Gfral-expressing Neurons Suppress Food Intake via Aversive Pathways"

Supplemental Figure 1

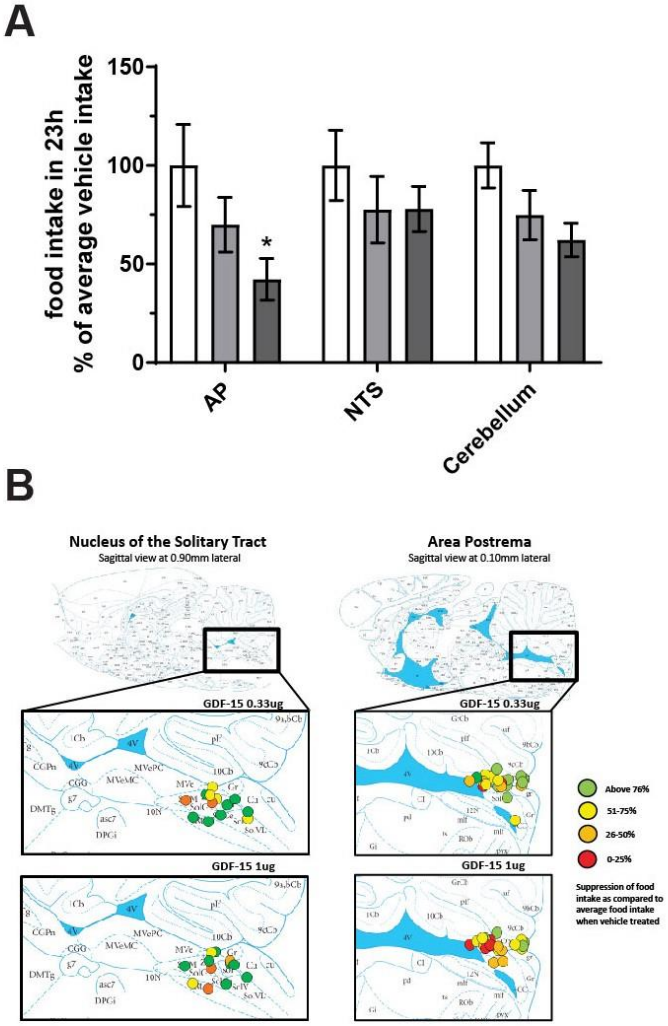

Supplemental Figure 2

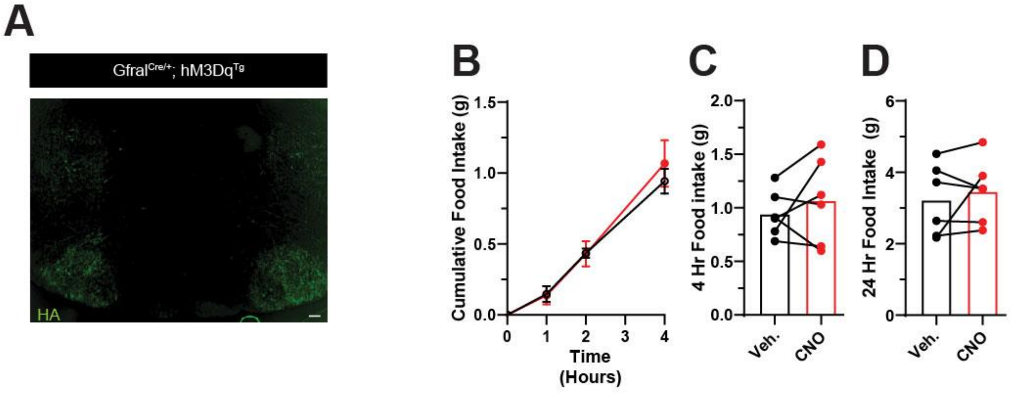

Supplemental Figure 3

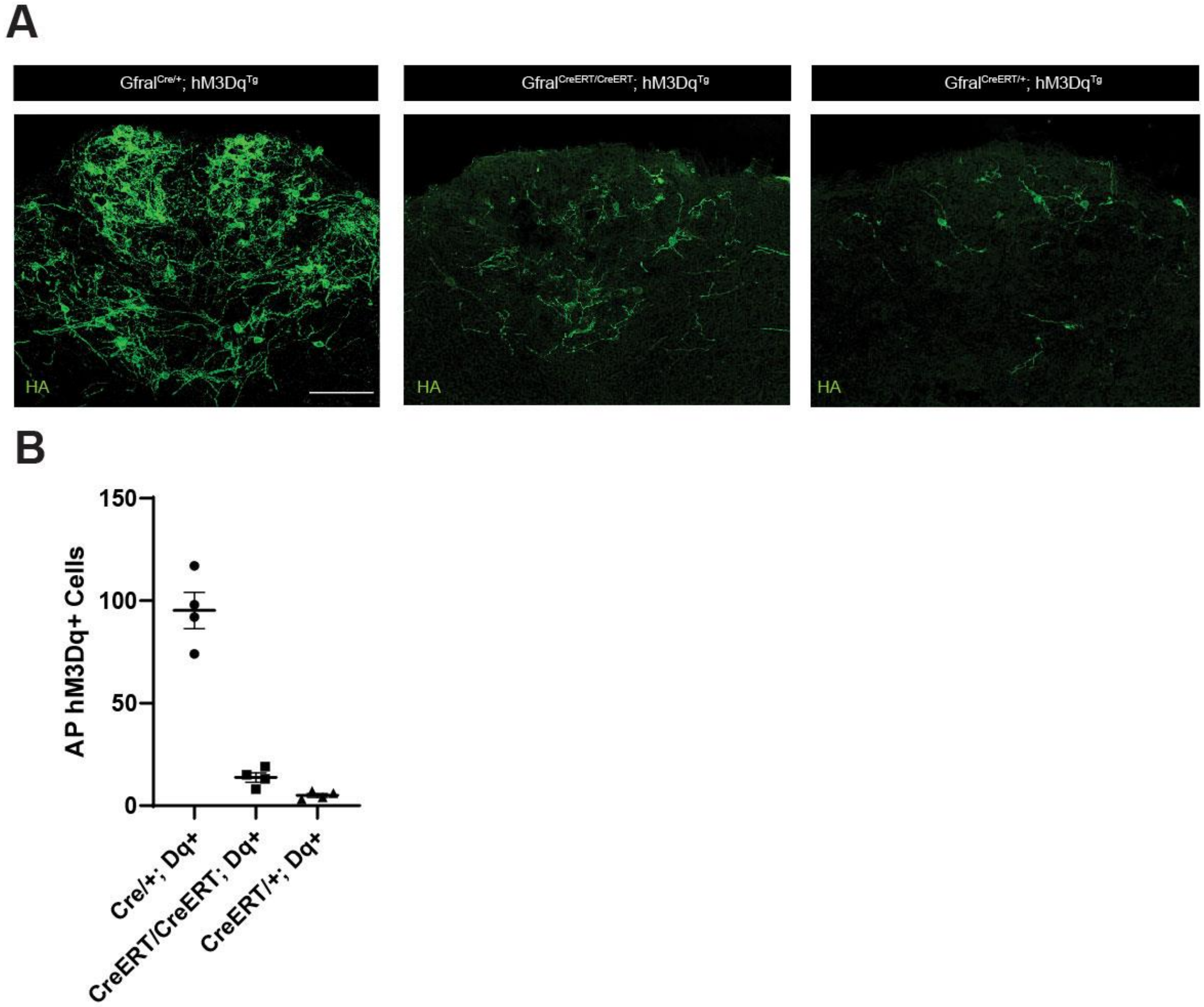

Supplemental Figure 4

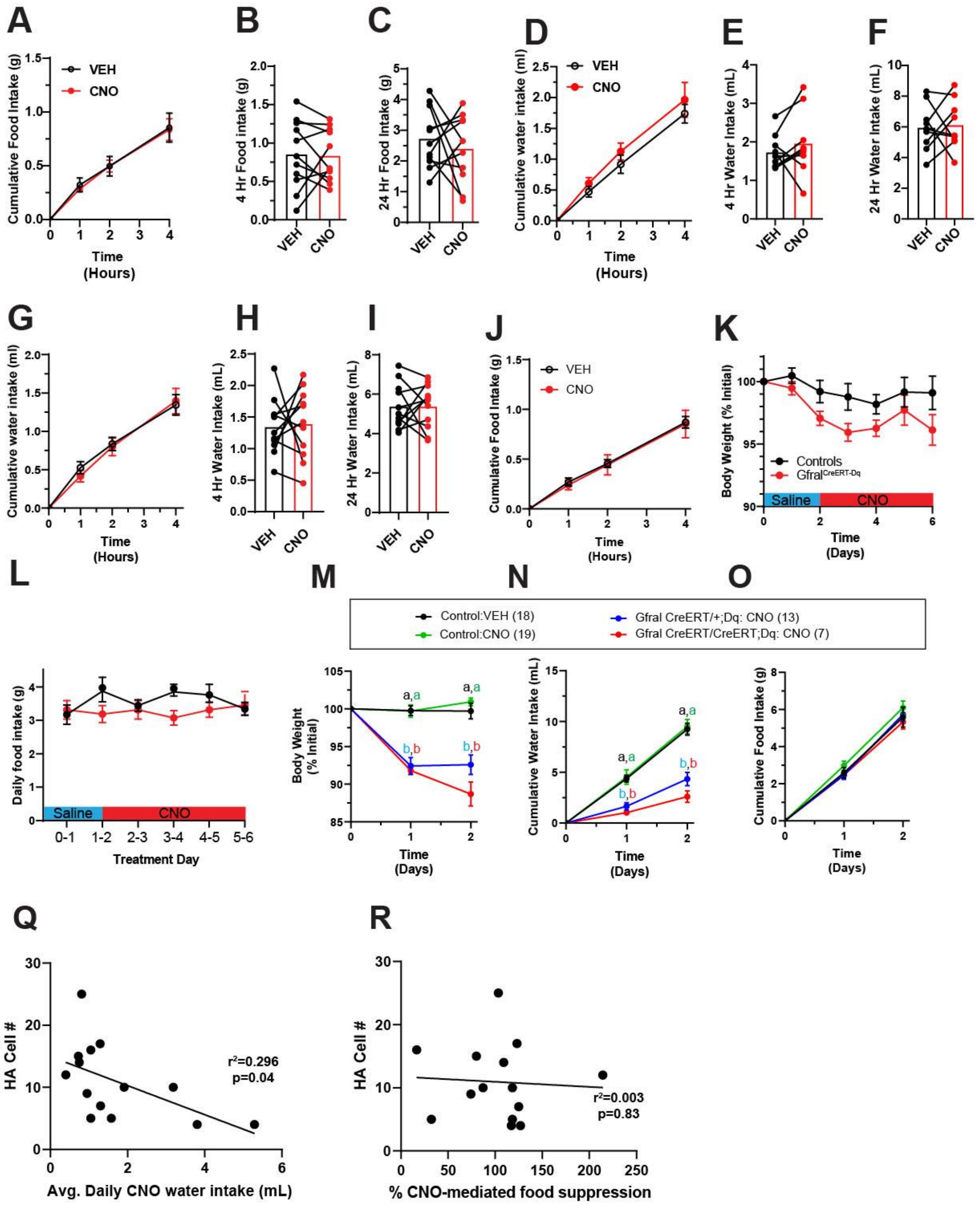

**Supplemental Figure 1: GDF-15 acts in the Area Postrema to reduce food intake.** (A) 23-h food intake in wildtype rats following infusion of either vehicle, 0.33 μg GDF-15 or 1 μg GDF-15 (data presented as percent of average vehicle intake). (B) "Anatomical heatmaps" with each circle representing infusion site within each rat. The colors represent the food the level of food suppression with green = above 76%, orange = 51-75%, yellow = 26-50%, and red = 0-25% of average vehicle food intake in all rats (n=8-14). All bars are represented as mean±SEM and * = $p<0.05$ as compared to vehicle intake in Dunnett's multiple comparison post hoc test following Two-way ANOVA. Scale bar =100 μm

**Supplemental Figure 2: DREADD expression in GFRAL$^{Cre\text{-}Dq}$ mice and controls for GFRAL$^{Cre\text{-}Dq}$ experiments**. (A) hM3Dq DREADD expression (green) within the facial nucleus of GFRAL$^{Cre\text{-}Dq}$ mice. (B-D) Dark phase food intake in control animals (lacking either DREADD or Cre expression) injected with vehicle (VEH, black) or CNO (red) (n=6). All bars are represented as mean±SEM

**Supplemental Figure 3: Number of DREADD expressing cells across *Gfral*$^{Cre}$ models.** (A) Representative images of AP staining of DREADD (HA, green) in GFRAL$^{Cre\text{-}Dq}$ (left panel), Tamoxifen(TMX)-treated GFRAL$^{CreERT\text{-}Dq}$ (*Gfral*$^{CreERT/CreERT}$ homozygotes)(center panel) and TMX-treated GFRAL$^{CreERT\text{-}Dq}$ (*Gfral*$^{CreERT/+}$ heterozygotes) (right panel). (B) Quantification of HA-expressing cells within the AP of each model (n=4). All data are presented as mean±SEM. Scale bar= 100 μm

**Supplemental Figure 4**: **Food and water intake studies for GFRAL$^{CreERT-Dq}$ and Controls.** (A-C) Dark phase food intake in controls for GFRAL$^{CreERT-Dq}$ experiments (n=11). (D-F) Dark phase water intake (no prior water deprivation) in GFRAL$^{CreERT-Dq}$ animals (n=11). (G-I) Dark phase water intake (no prior water deprivation) in controls administered CNO or vehicle 30 minutes prior to the onset of the dark cycle (n=11). (J) Food intake of DIO GFRAL$^{CreERT-Dq}$ animals administered CNO or vehicle (n=4). (K-L) Body weight and daily food intake of GFRAL$^{CreERT-Dq}$ mice administered with twice daily doses of saline for 2 days followed by twice daily doses of CNO for 4 days (n=7-9). (M-O) Water intake, body weight and food intake of mice with a single copy CreERT (*Gfral* $^{CreERT/+}$;Dq), two copies of CreERT (Gfral $^{CreERT/CreERT}$;Dq) mice and control mice exposed to control drinking water (Veh) or CNO-laced drinking water (CNO) (n values in legend). (Q-R) Linear regression analysis comparing number of HA-tagged DREADD expressing AP neurons to either the two-day average of CNO-containing drinking water intake or CNO-mediated food suppression (4 hours post onset of the dark cycle). All bars are represented as mean±SEM Two-way ANOVA with Dunnett's multiple comparison post hoc test was performed.
